# A CD4-CD8 T-cell circuit converts cardiac inflammation into tissue injury

**DOI:** 10.64898/2026.09.18.751575

**Authors:** Amir Z. Munir, Alan Gutierrez, Cade J. Krawiec, Riad Ghandour, Richard A. Baylis, Anya C. Shyani, Eva Rodriguez, Enrique Ortega-Sollero, Aishwarya Nene, Javid J. Moslehi

**Affiliations:** Section of Cardio-Oncology & Immunology, Cardiovascular Research Institute (CVRI), University of California San Francisco, School of Medicine, San Francisco, CA, USA; Yale University School of Medicine, New Haven, CT, USA; Spanish National Cardiovascular Research Centre (CNIC), Madrid, Spain; Department of Medicine, University of California San Francisco, School of Medicine, San Francisco, CA, USA

## Abstract

Inflammatory T-cell infiltration is widely considered a hallmark of cardiac immune-mediated tissue injury, yet whether inflammation alone is sufficient to cause cardiac damage remains unclear. Using a spontaneous genetic model of immune checkpoint inhibitor myocarditis, we show that cardiac immune infiltration and tissue injury are separable processes. CD4^+^ T-cells promote disease by licensing pathogenic CD8^+^ T-cell responses through CD40L signaling, whereas perforin-dependent CD8^+^ cytotoxicity is specifically required to induce cardiomyocyte death and cardiac arrhythmias. Loss of perforin prevented cardiac injury and rescued survival despite persistent myocardial inflammation, demonstrating that inflammatory infiltration is insufficient to produce lethal myocarditis in the absence of a cytotoxic effector program. CD40L blockade similarly attenuated pathogenic CD8^+^ T-cell activation, myocardial inflammation, and mortality. These findings identify a cooperative CD4–CD8 T-cell circuit that governs autoimmune cardiac injury and establish that acquisition of cytotoxic effector function, rather than inflammation alone, determines the transition from immune infiltration to tissue destruction.

## Introduction

T-cell infiltration is a hallmark of cardiac immune-mediated tissue injury, but inflammation does not invariably result in organ damage. What determines whether an inflammatory T-cell response progresses to tissue destruction remains poorly understood. In myocarditis, inflammatory T-cell infiltration and cardiomyocyte death are defining pathological features and form the basis of histologic diagnosis. However, whether infiltrating T-cells are themselves sufficient to drive cardiomyocyte injury, or whether tissue destruction requires acquisition of a specialized cytotoxic effector program, remains unknown.

Immune checkpoint inhibitor (ICI)-associated myocarditis provides a tractable system to address this question. ICIs enhance anti-tumor immunity by blocking inhibitory receptors such as PD-1 and LAG-3 but can also unleash autoreactive T-cell responses against healthy tissues. Although uncommon, ICI-myocarditis is among the most lethal immune-related adverse events, with mortality approaching 50% despite intensive immunosuppression(1). Histologically, the disease is characterized by dense myocardial T-cell infiltration and cardiomyocyte death, yet the cellular interactions that couple these processes remain incompletely understood(2). While CD8^+^ T-cells are generally viewed as the principal effectors, little is known about the role of CD4^+^ T-cells, raising the question of how these populations may cooperate to produce pathogenic cardiac responses. Other forms of autoimmune myocarditis have been linked to distinct CD4^+^ T-cell programs, but whether similar mechanisms operate in ICI-myocarditis is unknown(3, 4).

To address this question, we used a spontaneous genetic model of ICI-myocarditis in mice generated by combined deletion of *Lag3* and *Pdcd1* (genes encoding LAG-3 and PD-1, respectively), which faithfully recapitulates the cardiac inflammation, arrhythmias, and premature mortality observed in patients(5). We identify a cooperative T-cell circuit that separates myocardial inflammation from tissue injury. CD4^+^ T-cell and CD40L signaling sustain pathogenic CD8^+^ T-cell responses, whereas perforin-dependent cytotoxicity determines whether myocardial inflammation progresses to cardiomyocyte injury, arrhythmias, and death.

## Results

### Both CD4^+^ and CD8^+^ T-cells are necessary for lethality in ICI-myocarditis

To characterize the T-cell subsets in the *Lag3^-/-^,Pdcd1^-/-^* model of myocarditis, we performed flow cytometry and immunofluorescence which revealed an expansion of both cardiac CD4^+^ and CD8^+^ T-cells (**Figure 1A-B**). To test the necessity of both CD4^+^ and CD8^+^ T-cells, mice were injected with depleting CD4, CD8, or isotype control antibodies from 3 to 10 weeks of age **(Figure 1C)**. Effective depletion was confirmed by flow cytometry analysis of the cardiac T-cells (**Figure 1D**). Although cardiac CD8^+^ T-cells greatly outnumbered CD4^+^ T-cells, depletion of either subset completely rescued premature lethality **(Figure 1E)**, demonstrating that both populations are required for fulminant myocarditis. Electrographic analyses demonstrated that, despite the rescue of survival, the majority of anti-CD4 treated mice still developed arrhythmias which were absent in the anti-CD8 treatment suggesting that depletion of CD8 resulted in a more robust improvement (**Figure 1F**). Cardiac histology identified pronounced inflammation in isotype treated mice, mild inflammation in the anti-CD4 cohort, and essentially no inflammation in the anti-CD8 cohort (**Figure 1G-H**). Taken together, these results highlight that both CD4^+^ and CD8^+^ T-cells are necessary to induce fulminant myocarditis in the *Lag3^-/-^,Pdcd1^-/-^* model.

**Figure 1.**
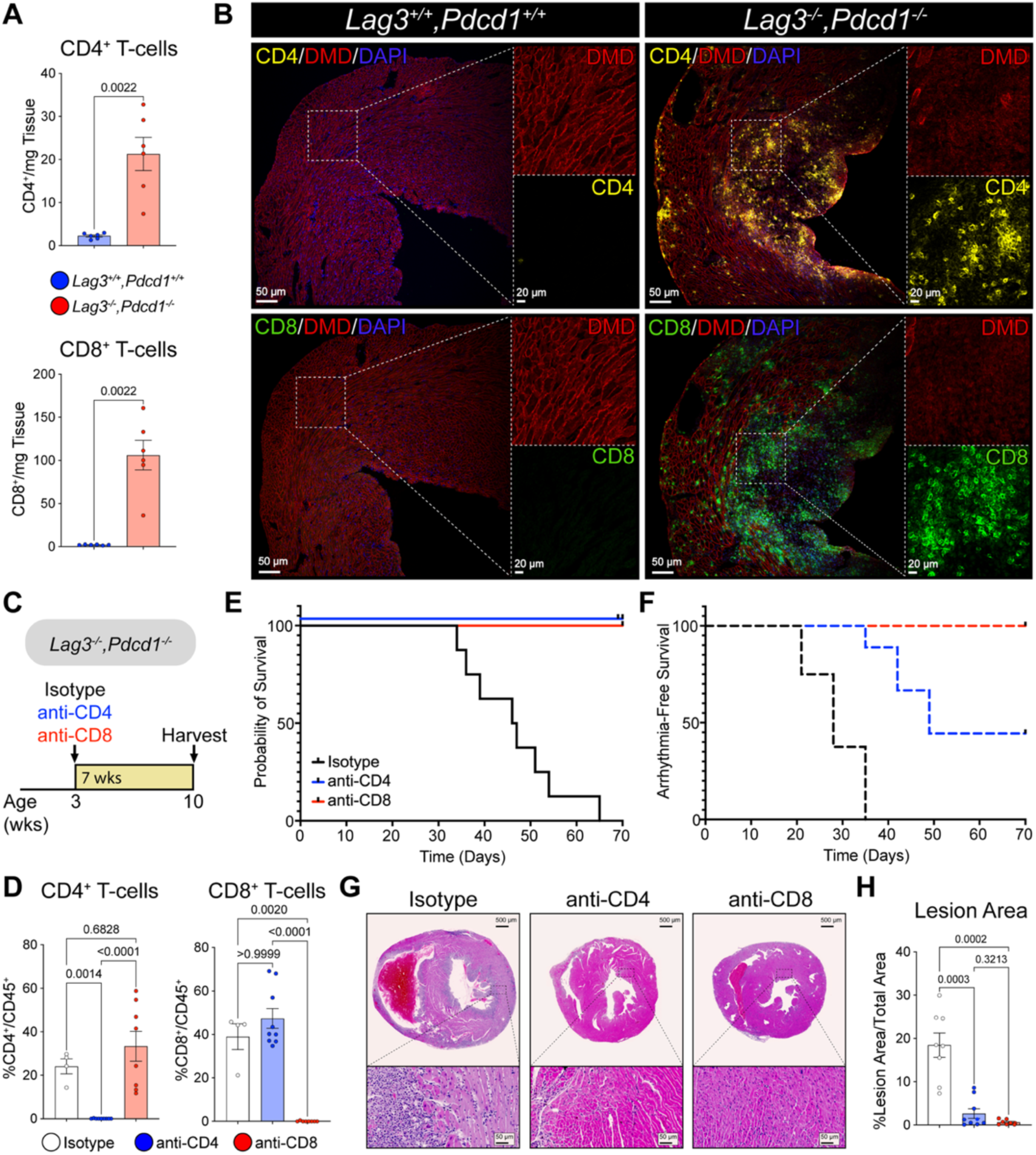
CD4^+^ and CD8^+^ T-cells are necessary for lethality in *Lag3^-/-^,Pdcd1^-/-^* myocarditis. **A.** Flow cytometry of cardiac CD4^+^ and CD8^+^ T-cell populations in *Lag3^-/-^,Pdcd1^-/-^* and control *Lag3^+/+^,Pdcd1^+/+^* mice. **B.** Immunofluorescence of CD4^+^ and CD8^+^ T-cells and Dystrophin (DMD) protein in *Lag3^-/-^,Pdcd1^-/-^*and control *Lag3^+/+^,Pdcd1^+/+^* mice. **C**. Diagram of treatment of *Lag3^-/-^,Pdcd1^-/-^* mice with anti-CD4, anti-CD8, and Isotype. **D.** Flow cytometry of cardiac CD4^+^ and CD8^+^ T-cell populations in Isotype, anti-CD4 and anti-CD8 treatment. **E**. Survival curve of Isotype, anti-CD4 or anti-CD8 treatment in *Lag3^-/-^,Pdcd1^-/-^* mice**. F.** Arrhythmia-free survival curve of Isotype, anti-CD4 or anti-CD8 treatment in *Lag3^-/-^,Pdcd1^-/-^* mice**. G.** Representative cardiac H&E histology of Isotype, anti-CD4 or anti-CD8 treatment in *Lag3^-/-^,Pdcd1^-/-^* mice. **H.** Quantification of myocarditis lesion area of Isotype, anti-CD4 or anti-CD8 treatment in *Lag3^-/-^,Pdcd1^-/-^*mice.

### CD4^+^ T-cells in *Lag3^-/-^,Pdcd1^-/-^* myocarditis are enriched in *Gzmk* and Th1 signatures

The most widely used pre-clinical model of autoimmune myocarditis involves immunizing BALB/c mice with an alpha myosin peptide and has been strongly linked to Th17 biology(3, 6). In contrast, prior models of ICI-myocarditis have been predominantly CD8 mediated(7–9). The CD4^+^ T-cell contribution to ICI-myocarditis is poorly understood. Given that CD4 depletion rescued survival, we sought to characterize the relevant cardiac CD4^+^ T-cell populations in the *Lag3^-/-^,Pdcd1^-/-^* mice. Single cell RNA sequencing on sorted cardiac CD4^+^ T-cells from *Lag3^-/-^,Pdcd1^-/-^* and wild-type (WT) mice identified naïve, proinflammatory T helper 1 (Th1), regulatory T-cell (Treg), *Granzyme K* (*Gzmk*) expressing, and dividing CD4^+^ T-cell clusters (**Figure 2A-B**). Among these CD4 clusters, the Th1 and *Gzmk* clusters were the most expanded in myocarditis (**Figure 2C**). We found the *Gzmk* expressing cluster to be of particular interest, as granzymes are canonically thought to play roles in cell-mediated cytotoxicity, with recent evidence suggesting *Gzmk* may act as a complement activator with important roles in autoimmune disease(10, 11). Therefore, we hypothesized that loss of *Gzmk* expression in CD4^+^ T-cells may attenuate myocarditis. We performed inducible CD4-specific knockout of Gzmk with *Cd4-*Cre^ERT2^ *Gzmk^fl/fl^,Lag3^-/-^,Pdcd1^-/-^* mice treated with tamoxifen chow from 3 to 10 weeks of age, which showed evidence of cre-mediated excision (**Figure 2D; Figure S1)**. Contrary to our hypothesis, loss of Gzmk in CD4^+^ T-cells had no discernible reduction in the severity of myocarditis with similar premature lethality (**Figure 2E**), This was further supported by a similar degree of myocardial inflammation on histology (**Figure 2F-G**). In summary, loss of *Gzmk* in CD4^+^ T-cells did not alter the cardiac phenotype of our preclinical ICI myocarditis model.

**Figure 2.**
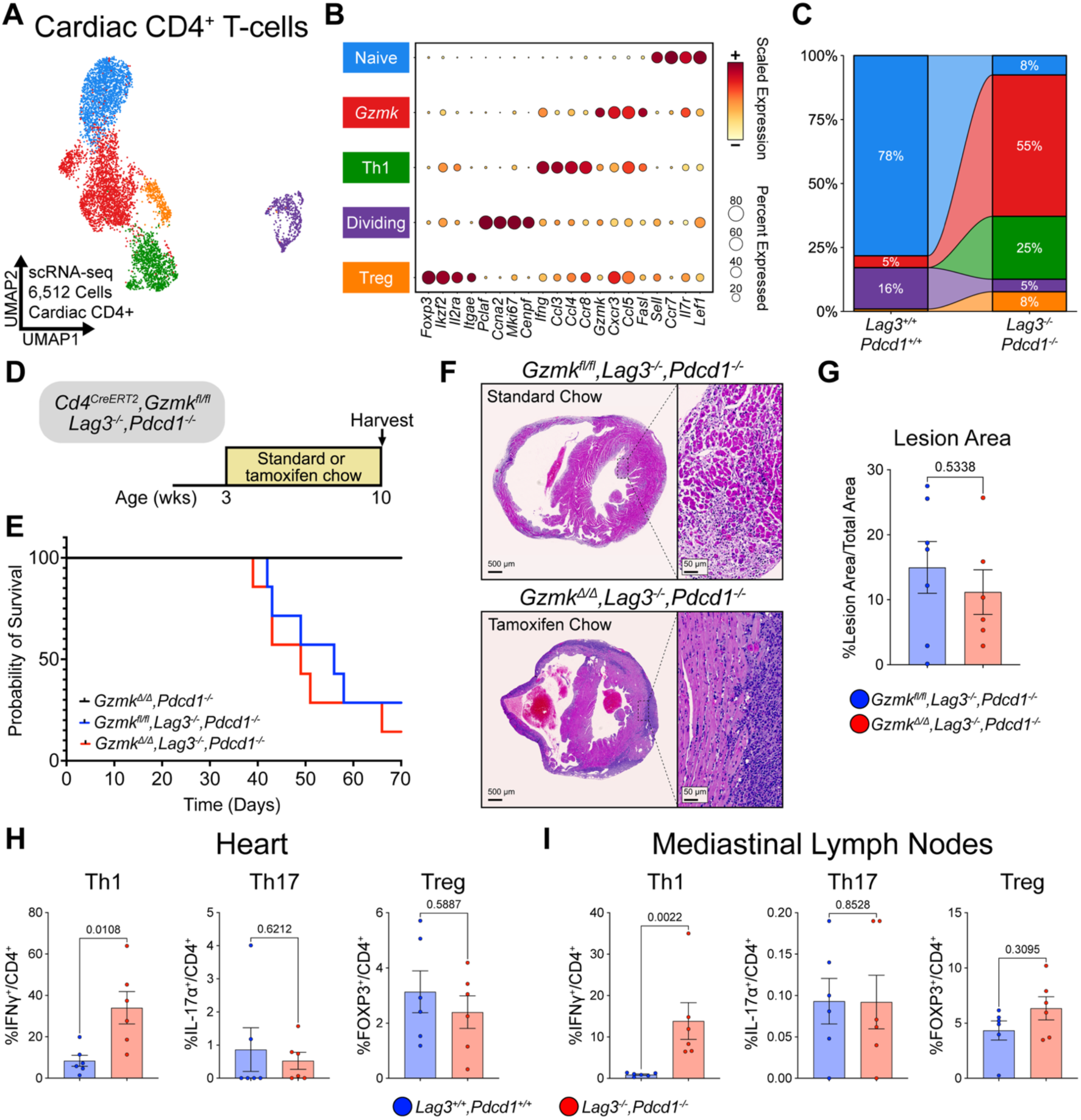
Cardiac CD4^+^ T-cells in *Lag3^-/-^,Pdcd1^-/-^* myocarditis are enriched in *Gzmk* and Th1 signatures. **A.** UMAP of Cardiac CD4^+^ sorted scRNA-seq from 4 *Lag3^-/-^,Pdcd1^-/-^* and 8 *Lag3^+/+^,Pdcd1^+/+^* mice. **B.** DotPlot of marker genes of each cardiac CD4^+^ scRNA-seq clustering. **C.** Stacked barplot of the representative CD4^+^ populations in *Lag3^-/-^,Pdcd1^-/-^* and *Lag3^+/+^,Pdcd1^+/+^*mice. **D.** Diagram of genetic mouse model of inducible loss of *Gzmk* in CD4 expressing T-cells on a *Lag3^-/-^,Pdcd1^-/-^* background. **E.** Survival curve of tamoxifen-fed *Cd4-*Cre^ERT2^ *Gzmk^fl/fl^, Lag3^+/+^, Pdcd1^-/-^,* regular chow-fed and tamoxifen-fed *Cd4-*Cre^ERT2^ *Gzmk^fl/fl^, Lag3^-/-^, Pdcd1^-/-^*. **F.** Representative H&E histology of regular chow-fed and tamoxifen-fed *Cd4-*Cre^ERT2^ *Gzmk^fl/fl^, Lag3^-/-^, Pdcd1^-/-^*mice. **G.** Quantification of myocarditis lesion area in regular chow-fed and tamoxifen-fed *Cd4-*Cre^ERT2^ *Gzmk^fl/fl^, Lag3^-/-^, Pdcd1^-/-^* mice. **H.** Cardiac CD4^+^ helper T-cell subset flow in *Lag3^-/-^,Pdcd1^-/-^*and *Lag3^+/+^,Pdcd1^+/+^* mice. **I.** Mediastinal lymph node CD4^+^ helper T-cell subset flow in *Lag3^-/-^,Pdcd1^-/-^*and *Lag3^+/+^,Pdcd1^+/+^* mice.

### CD8^+^ T-cell responses in ICI-myocarditis are dependent on CD40L signaling

We started by confirming the Th1 predominance in our *Lag3^-/-^,Pdcd1^-/-^*mice by performing flow cytometry of cardiac tissue and mediastinal lymph nodes. Consistent with the scRNA-seq data, Th1 CD4^+^ T-cells expanded in myocarditis with no changes in the Th17 population (**Figure 2H-I**). Similar results were observed in the spleen and blood (**Figure S2**).

Given the dominant Th1 cardiac inflammation in our model, we hypothesized that Th1 T-cells potentiated CD8^+^ T-cell propagation during myocarditis. Previous literature has highlighted the role of CD40L, a costimulatory molecule expressed on CD4^+^ T-cells, in priming CD8^+^ T-cell-mediated clearance during infection and in anti-tumor immunity(12–15). Consistent with this hypothesis, CD40L expression was higher in the CD4^+^ T-cells in the mediastinal lymph nodes of *Lag3^-/-^,Pdcd1^-/-^* mice as compared to WT controls (**Figure 3A**). We therefore sought to determine the necessity of CD40L signaling in fulminant myocarditis using our model (**Figure 3B**). Treatment of 3-week-old *Lag3^-/-^,Pdcd1^-/-^* mice with neutralizing anti-CD40L antibody or isotype control for a total of 7 weeks rescued mortality with 75% of CD40L antibody-treated mice surviving while all the isotype-treated mice had died by 10 weeks of age (**Figure 3C**). In line with the improvement in survival anti-CD40L treated mice experienced fewer arrhythmias (**Figure 3C**) and had a significant reduction in myocardial inflammation on histology (**Figure 3D-E**). To determine the changes in the immune repertoire with CD40L neutralization, we performed flow cytometry on the heart and mediastinal lymph nodes. Treatment with anti-CD40L for 7-weeks resulted in decreased cardiac T-cell, CD8^+^ T-cell, and CD8^+^ effector memory T-cells (CD8^+^ T_EM_), with similar trends identified in the 3-week treatment (**Figure 3F**). Assessment of mediastinal lymph node CD4^+^ T_EM_ and CD8^+^ T_EM_ populations showed significant reductions at both 3 and 7-weeks of anti-CD40L treatment (**Figure 3G**). To further phenotype the cardiac immune populations, we performed CD45^+^ sorted scRNA-seq of anti-CD40L or isotype treated mice after 3-weeks of treatment. Again, we were able to identify a dynamic reduction in activated T-cell clusters (*Cxcr3/Gzmk* and *Cd8/Prf1* T-cell clusters) with CD40L neutralization (**Figure 3H**). Further T-cell sub-clustering revealed a relative expansion of naïve cardiac T-cells with CD40L blockade and reduction in activated CD8^+^ T-cells expressing *Prf1*, *Ly6c2,* and *Gzmk* (**Figure 3I**). Taken together, these data show that CD4^+^ T-cells support pathogenic CD8^+^ T-cell expansion through CD40L signaling.

**Figure 3.**
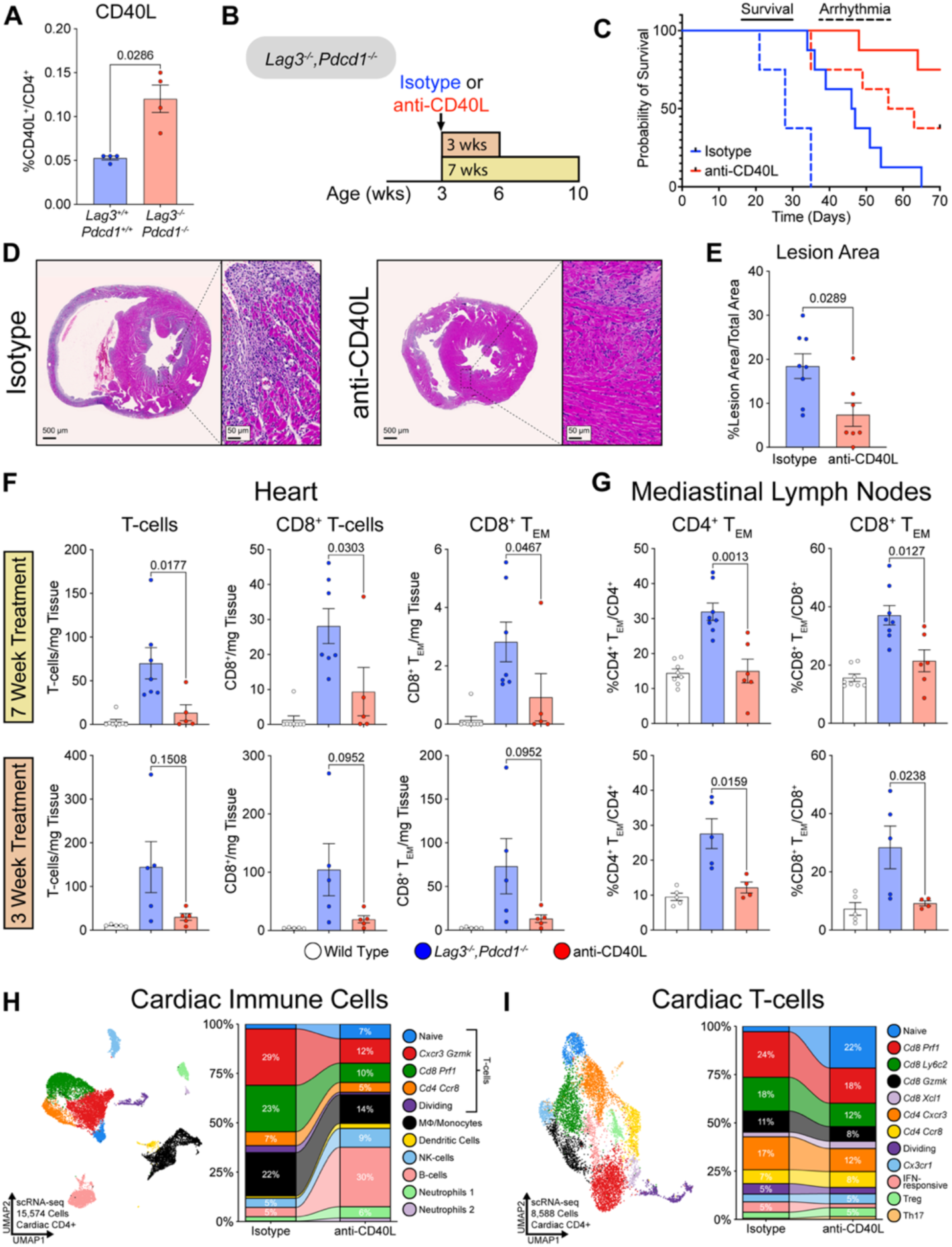
CD8^+^ T-cell responses in *Lag3^-/-^,Pdcd1^-/-^* myocarditis are dependent on CD40L signaling. **A.** Mediastinal lymph node CD40L flow cytometry in *Lag3^-/-^,Pdcd1^-/-^*and *Lag3^+/+^,Pdcd1^+/+^* mice. **B.** Diagram of isotype vs anti-CD40L treatment of *Lag3^-/-^,Pdcd1^-/-^* mice. **C.** Survival and Arrhythmia-free survival curve of 7-week isotype vs anti-CD40L treatment. **D.** Representative cardiac H&E histology of 7-week isotype vs anti-CD40L treatment. **E.** Quantification of myocarditis lesion area in 7-week isotype vs anti-CD40L treatment. **F.** Flow Cytometry of cardiac T-cell, CD8^+^ T-cell, and effector memory CD8^+^ T-cell (CD8^+^ T) in 3-week and 7-week isotype vs anti-CD40L treatment. **G.** Flow Cytometry of mediastinal lymph node CD4^+^ and CD8^+^ effector memory in 3-week and 7-week isotype vs anti-CD40L treatment. **H.** UMAP and stacked bar plot of cardiac CD45^+^ sorted scRNA-seq of 3-week isotype vs anti-CD40L treatment. **I.** UMAP and stacked bar plot of cardiac T-cell subsets of 3-week isotype vs anti-CD40L treatment.

### Perforin-1 is essential for CD8^+^ T-cell mediated cardiac injury in ICI-myocarditis

Because our data suggested that CD4^+^ T-cells primarily amplify rather than directly mediate cardiomyocyte injury, we next examined cytotoxic pathways in CD8^+^ T-cells. CD8^+^ T-cells can mediate target-cell death through Fas-Fas-ligand (FasL) signaling or through the perforin-granzyme pathway, in which perforin-1 forms pores that permit delivery of granzyme B (16, 17). FasL neutralization had no effect on premature lethality or arrhythmia burden in *Lag3^-/-^,Pdcd1^-/-^* mice (**Figure S3**). In contrast, genetic deletion of Perforin-1 (*Prf1*) in *Lag3^-/-^,Pdcd1^-/-^* mice completely rescued survival and prevented arrhythmias by 10 weeks of age (**Figure 4A**). Despite this marked phenotypic rescue, *Prf1*^-/-^,*Lag3^-/-^,Pdcd1^-/-^*mice still showed cardiac inflammation on histology, though at lower levels compared to *Lag3^-/-^,Pdcd1^-/-^* mice (**Figure 4B-C**). These findings were corroborated by flow cytometry, which confirmed persistent cardiac immune infiltration, with elevated CD45^+^ leukocytes, CD4^+^ T-cells, and CD8^+^ T-cells in *Prf1*^-/-^, *Lag3^-/-^,Pdcd1^-/-^* mice compared to WT controls, although reduced relative to *Lag3^-/-^ ,Pdcd1^-/-^* mice (**Figure 4D**). Immunofluorescence similarly demonstrated persistent CD8^+^ T-cell infiltration within the myocardium in *Prf1*^-/-^,*Lag3^-/-^,Pdcd1^-/-^*mice (**Figure 4E**). Strikingly, perforin deficiency nearly abolished cardiomyocyte apoptosis and troponin release despite persistent inflammatory infiltrates: cleaved caspase-3 staining was minimal in Prf1-deficient *Lag3^-/-^,Pdcd1^-/-^* mice, and serum cardiac troponin I, a biomarker of cardiomyocyte damage, was markedly reduced **(Figure 4F-H).** In summary, these findings identify perforin-1 as the principal mediator of cardiomyocyte injury and demonstrate that loss of *Prf1* uncouples cardiac inflammation from injury in CD8-mediated myocarditis.

**Figure 4.**
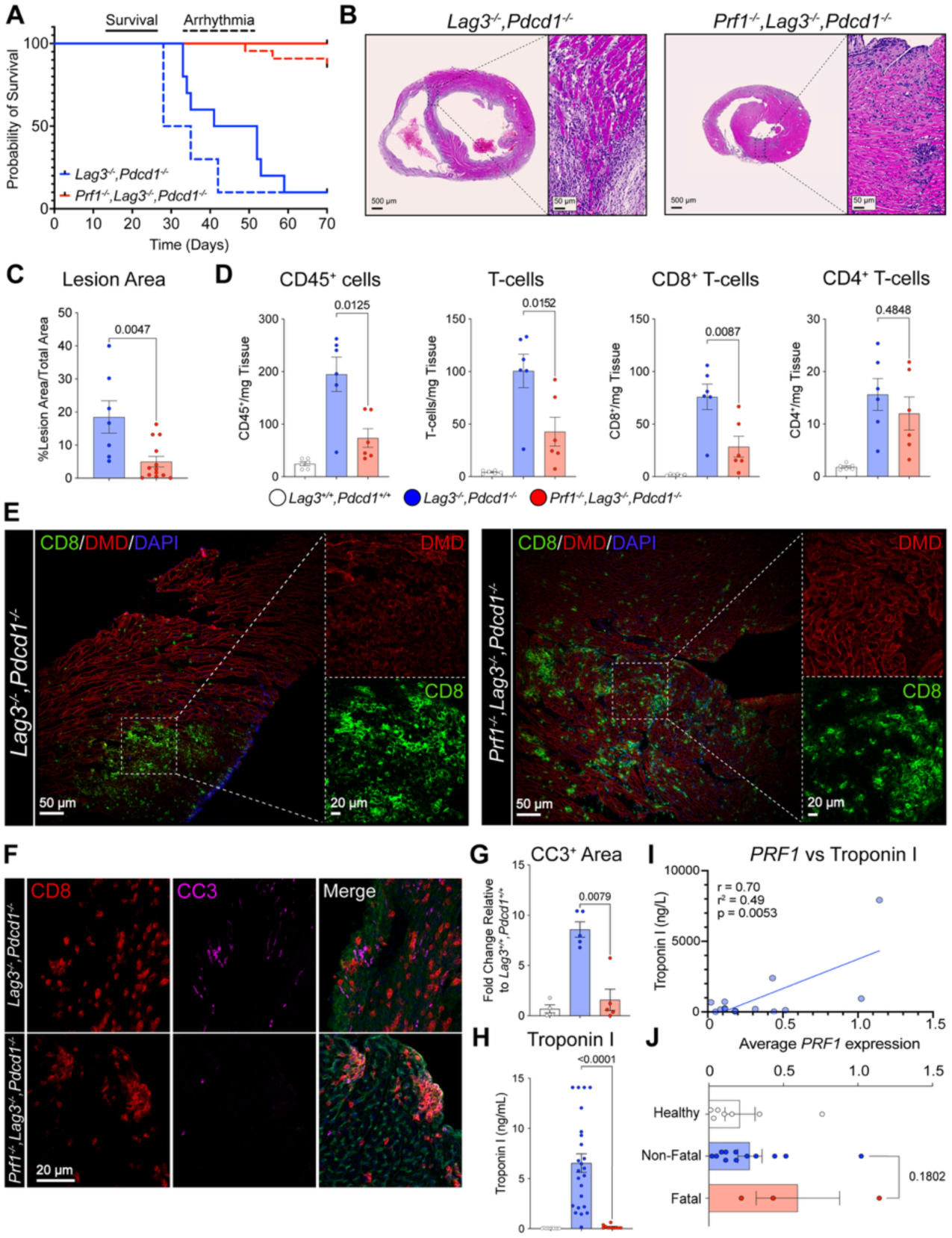
Perforin-1 is essential for CD8^+^ T-cell mediated cardiac injury in *Lag3^-/-^,Pdcd1^-/-^* mice. **A.** Survival and arrhythmia-free survival curves of *Lag3^-/-^,Pdcd1^-/-^*and *Prf1^-/-^,Lag3^-/-^,Pdcd1^-/-^* mice. **B.** Representative cardiac H&E histology of *Lag3^-/-^,Pdcd1^-/-^*and *Prf1^-/-^,Lag3^-/-^,Pdcd1^-/-^* mice. **C.** Quantification of myocarditis lesion area in *Lag3^-/-^,Pdcd1^-/-^* and *Prf1^-/-^,Lag3^-/-^,Pdcd1^-/-^* mice. **D.** Flow cytometry of CD45^+^, T-cells, CD8^+^ T-cells, and CD4^+^ T-cells in *Lag3^+/+^,Pdcd1^+/+^, Lag3^-/-^,Pdcd1^-/-^*and *Prf1^-/-^,Lag3^-/-^,Pdcd1^-/-^*mice. **E.** Representative CD8^+^ and DMD immunofluorescence of *Lag3^-/-^,Pdcd1^-/-^*and *Prf1^-/-^,Lag3^-/-^,Pdcd1^-/-^* mice. **F.** Representative CD8^+^ and cleaved caspase-3 (CC3) immunofluorescence of *Lag3^-/-^,Pdcd1^-/-^* and *Prf1^-/-^,Lag3^-/-^,Pdcd1^-/-^*mice. **G.** Quantification of whole slide CC3 staining in *Lag3^+/+^,Pdcd1^+/+^, Lag3^-/-^,Pdcd1^-/-^* and *Prf1^-/-^,Lag3^-/-^,Pdcd1^-/-^* mice. **H.** Serum cardiac troponin I values in *Lag3^+/+^,Pdcd1^+/+^, Lag3^-/-^,Pdcd1^-/-^* and *Prf1^-/-^,Lag3^-/-^,Pdcd1^-/-^*mice. **I.** Human serum troponin I compared to average cell *PRF1* expression from publicly accessed human ICI-myocarditis scRNA-seq dataset. **J.** Average cell *PRF1* expression between healthy controls, non-fatal ICI myocarditis and fatal ICI-myocarditis cases from publicly accessed human ICI-myocarditis scRNA-seq dataset.

To validate our pre-clinical findings, we turned to an existing dataset of cardiac scRNA-seq from ICI-treated patients and controls(18). In patients with ICI-myocarditis, average cardiac *Prf1* expression correlated with serum troponin levels **(Figure 4I)**. Further, average cardiac *Prf1* expression trended upwards in patients with fatal ICI-myocarditis cases as compared to non-fatal or healthy controls **(Figure 4J)**.

## Discussion

In this study, we show that cooperation of CD4^+^ and CD8^+^ T-cells is necessary for ICI-myocarditis through CD40L and perforin-1, respectively. Though both CD4^+^ and CD8^+^ T-cells were required for the lethality, these data suggest that CD4^+^ T-cells primarily support the disease process through CD40L-CD40 costimulation which enables exuberant pathogenic CD8^+^ T-cell responses. In turn, CD8^+^ T-cells inflict cardiomyocyte death through perforin-1. The finding that perforin-1 deletion prevents cardiomyocyte injury and lethality despite persistent cardiac inflammation suggests that immune infiltration and tissue damage are separable processes in CD8-mediated-myocarditis.

Although ICI-myocarditis is often considered a CD8^+^ T-cell-mediated disease, our depletion studies demonstrate that CD4^+^ T-cells are critical to induce severe disease. However, there were notable differences in the benefit of CD4 and CD8 depletion with CD4 depletion improving survival but still developing arrhythmias and mild inflammation. In contrast, the CD8 depletion produced more complete protection from arrhythmia and histologic myocarditis. These findings suggest that CD4^+^ T-cells are necessary upstream disease amplifiers, while CD8^+^ T-cells are more proximal mediators of cardiac injury. Mechanistically, this appears to be mediated by CD40 signaling as disruption of this pathway reduced mortality, arrhythmias, histologic myocarditis, and cardiac CD8^+^ effector memory expansion. Together, these data support a model in which CD40L signaling links the Th1-skewed CD4^+^ compartment to the expansion or maintenance of pathogenic CD8^+^ effectors.

Although our data implicate CD40L as a critical disease pathway, they do not establish whether CD40L acts primarily through dendritic cells, macrophages, B cells or another CD40-expressing population(19). Given the known role of CD40–CD40L interactions in licensing antigen-presenting cells (APCs) and promoting CD8^+^ T-cell immunity, one attractive model is that CD4^+^ T-cell CD40L enhances APC activation, thereby sustaining cytotoxic CD8^+^ T-cell responses in both the cardiac-draining lymph node and heart.

A second important finding is that, in contrast to the classic experimental autoimmune myocarditis (EAM) model, the cardiac CD4^+^ T-cell response in this model is dominated by Th1-associated inflammation rather than Th17(3, 6). These data support that myocarditis arising from checkpoint blockade may follow a distinct immunologic trajectory from EAM. The diversity of dominant T-cell phenotypes among myocarditis models raises the question of the utility of the term “lymphocytic myocarditis” which likely encompasses many different forms of cardiac inflammatory diseases that may depend on the original inciting trigger and immune substrate of the patient.

One of the more striking observations in this study is that *Prf1* deletion nearly abolishes lethality, arrhythmias, troponin elevation, and cardiomyocyte apoptosis, despite persistent cardiac immune infiltration. This indicates that cardiac inflammation alone is insufficient to produce the lethal phenotype in this model. Instead, perforin-dependent cytotoxicity appears to be a required effector mechanism through which CD8^+^ T-cells convert immune infiltration into cardiomyocyte injury. These data provide direct genetic evidence that cytotoxic granule release is a necessary mediator of myocardial damage. Analysis of human cardiac single-cell RNA-seq data provides supportive translational evidence for this mechanism. The association between cardiac PRF1 expression and serum troponin, together with the trend toward higher PRF1 expression in fatal ICI-myocarditis, suggests that perforin-associated cytotoxicity may also mark clinically severe disease in patients. However, these human data are correlative and derived from limited samples, so they should be interpreted as hypothesis-supporting rather than definitive evidence of causality.

These findings have several broader implications. First, they suggest that the severity of ICI-myocarditis may be determined not only by the magnitude of cardiac immune infiltration but by whether infiltrating T-cells acquire a cytotoxic program capable of directly injuring cardiomyocytes. As a corollary to this, our data raise the possibility that biomarkers of cytotoxicity, including PRF1-associated transcriptional programs, may better reflect tissue injury than inflammatory burden alone. Finally, the CD40L-CD8-Prf1 axis provides a framework for therapeutic strategies aimed at interrupting pathogenic T-cell help or cytotoxic effector function.

This study has several limitations. The *Lag3^-/-^,Pdcd1^-/-^* model reproduces key features of ICI-myocarditis but does not fully capture the timing or tumor context of human ICI exposure. In addition, *Prf1* was deleted globally, so future studies will be important to confirm the precise cellular source of pathogenic perforin. Similarly, although CD40L blockade strongly attenuated disease, these experiments do not identify the relevant CD40-expressing cellular target.

In summary, our findings define a CD4–CD8 T-cell circuit in which Th1-skewed CD4^+^ T-cells and CD40L signaling promote pathogenic CD8^+^ T-cell responses, while perforin-dependent cytotoxicity converts cardiac inflammation into cardiomyocyte injury, arrhythmias, and death. These results distinguish inflammatory infiltration from cytotoxic tissue damage and identify Prf1-mediated CD8^+^ T-cell injury as a central mechanism of ICI-myocarditis.

## Methods

### Sex as a biological variable

Both male and female mice were included across experiments. Sex was incorporated as a biological variable in experimental design; however, not every individual experiments was powered to detect sex-specific differences, and data from both sexes were therefore analyzed together.

### Animals

*Pdcd1* null*, Lag3* null, *Prf1, Cd4-*cre^ERT2^ null mice were purchased from the Jackson Laboratory (Strain 026644, 028276, 002407, 022356). *Gzmk*^fl/fl^ mice were obtained from David Oh, MD, PhD at UCSF. Lines were crossed to generate *Lag3^-/-^,Pdcd1^-/-^ ;Prf1^-/-^,Lag3^-/-^,Pdcd1^-/-^;* and *Cd4-*Cre^ERT2^ *Gzmk^fl/fl^,Lag3^-/-^,Pdcd1^-/-^* mice. Mice were housed at the UCSF Cardiovascular Research Institute Barrier, an animal facility accredited by the Association for Assessment and Accreditation of Laboratory Animal Care International. Animals were maintained in a controlled environment with a 12-hour light-dark cycle with access to water and a chow diet at all times. All experiments were performed in accordance with an IACUC protocol (AN207188). The study adheres to the ARRIVE guidelines for reporting animal research. Both male and female mice were studied in equal number. Mice were euthanized via deep isoflurane anesthesia followed by cervical dislocation. Mouse survival was assessed via Kaplan-Meier Survival curves with log rank statistical analysis.

### Generation of single cell cardiac suspension

Mice were euthanized and hearts were perfused via the left ventricle with 10 mL cold PBS, excised, and finely minced. Tissue was enzymatically digested using the Multi Tissue Dissociation Kit 2 (Miltenyi Biotec, 130-110-203) according to the manufacturer’s protocol, with enzyme concentrations adjusted to preserve immune cell epitope integrity and diluted in Roswell Park Memorial Institute (RPMI) medium. Samples were incubated at 37 °C for 50 minutes with shaking at 185 RPM. Digested tissue was passed through a 70 µm cell strainer and washed with PEB buffer. Debris was removed using the Debris Removal Solution (Miltenyi Biotec, 130-109-398) following the manufacturer’s protocol. Red blood cells were lysed with ACK lysing buffer (Fisher Scientific, NC0274127) for 2 minutes at room temperature, then quenched with FACS buffer. Cells were pelleted and resuspended in FACS buffer.

### Cardiac CD4^+^ T-cell scRNA-seq

Cells were stained with PerCP/Cyanine 5.5 anti-mouse CD45 (BioLegend, 103132, clone 30-F11) and APC anti-mouse CD4 (BioLegend, 100412, clone GK1.5) for 30 minutes at 4°C. For the final 5 minutes, cells were also stained with DAPI (1:10,000). Live CD45^+^ CD4^+^ immune cells were sorted by fluorescence-activated cell sorting on PerCP/Cyanine 5.5 positive, APC positive, DAPI negative events. Given the paucity of cardiac CD4^+^ T-cells at homeostasis, the *Lag3^+/+^,Pdcd1^+/+^* mice consisted of pooled cardiac immune infiltrates from eight mice. The myocarditis sample consisted of inflamed hearts from four *Lag3^-/-^,Pdcd1^-/-^* mice. All mice were approximately six weeks of age.

Each sample (targeting 5,000 to 15,000 cells per sample) was processed for single-cell 5’ RNA using the 10x Genomics Chromium Next GEM 5’ Gene Expression Reagents v2. Libraries were prepared following the manufacturer protocol with the assistance of Gladstone Genomics Core. The libraries were sequenced using NovaSeq X and analysis was completed using 10x Genomics Cell Ranger software. We identified 7,046 cells from pooled *Lag3^-/-^,Pdcd1^-/^*^-^ mice (1,750 median genes per cells; 69,718 reads per cell) and 5,042 cells from pooled *Lag3^+/+^,Pdcd1^+/+^* mice (1,216 median genes per cell; 55,518 reads per cell). Data were analyzed in R using the filtered h5 matrices in Seurat. In brief, samples were subset to include cells with >200 but <4000 unique transcripts to exclude probable noncellular RNA reads and doublets, respectively.

The DoubletFinder R package was used to detect and remove doublets from analysis. After removing contaminants, five clusters were identified using a resolution of 0.1. Uniform Manifold Approximation and Projection was used for dimensionality reduction with 20 nearest neighbors and minimum distance of 0.3. The FindMarkers Seurat function was used with minimum percentage of 20% and minimum log2fold change of 1.

### Cardiac CD45^+^ T-cell scRNA-seq of isotype and anti-CD40L treated mice

Cells were stained with PerCP/Cyanine 5.5 anti-mouse CD45 (BioLegend, 103132, clone 30-F11) for 30 minutes at 4°C. For the final 5 minutes, cells were also stained with DAPI (1:10,000). Live CD45^+^ immune cells were sorted by fluorescence-activated cell sorting on PerCP/Cyanine 5.5 positive, DAPI negative events. The isotype-treated *Lag3^-/-^,Pdcd1^-/-^* mice consisted of pooled cardiac immune infiltrates from two mice. The anti-CD40L-treated *Lag3^-/-^,Pdcd1^-/-^* mice consisted of pooled cardiac immune infiltrates from four mice. All mice were approximately six weeks of age.

Each sample (targeting 10,000 to 20,000 cells per sample) was processed for single-cell 5’ RNA using the 10x Genomics Chromium GEM-X 5’ Gene Expression Reagents v3. Libraries were prepared following the manufacturer protocol with the assistance of Gladstone Genomics Core. The libraries were sequenced using NovaSeq X and analysis was completed using 10x Genomics Cell Ranger software. We identified 16,830 cells from pooled isotype-treated *Lag3^-/-^,Pdcd1^-/-^* mice (3,816 median genes per cells; 13,296 reads per cell) and 7,181 cells from pooled anti-CD40L-treated *Lag3^-/-^,Pdcd1^-/-^* mice (3,307 median genes per cell; 10,286 reads per cell). Data were analyzed in R using the filtered h5 matrices in Seurat. In brief, samples were subset to include cells with >200 but <5000 unique transcripts to exclude probable noncellular RNA reads and doublets, respectively. The DoubletFinder R package was used to detect and remove doublets from analysis. After removing contaminants, eleven clusters were identified using a resolution of 0.15. Uniform Manifold Approximation and Projection was used for dimensionality reduction with 20 nearest neighbors and minimum distance of 0.3. The FindMarkers Seurat function was used with minimum percentage of 35% and minimum log2fold change of 1.

### Flow Cytometry

*Lag3-/-, Pdcd1-/-* samples were run on an Attune NxT Acoustic Focusing cytometer (Life Technologies). Data were collected using Attune NxT software v.3.2.1. Analysis was performed in FlowJo v.10.10. The gating strategy consisted of forward scatter and side scatter to exclude debris, FSC area versus FSC height to exclude doublets, and Zombie Violet Viability to exclude dead cells (VWR, 10761-308). Gating strategy can be found in **Figure S4**. The following antibodies were used for *Lag3-/-, Pdcd1-/-* samples: CD45-PerCP/Cy5.5 (BioLegend, 103132, clone 30-F11; dilution 1:400); CD3-AF488 (BioLegend, 100210, clone 17A2; dilution 1:200); CD4-APC (BioLegend, 100412, clone GK1.5; dilution 1:100); CD8a-PE/Cy7 (BioLegend, 100722, clone 53-6.7, dilution 1:400), TCR-Beta-PE/eFlour610 (Invitrogen, 61-5961-82, clone H57-597; dilution 1:100), CD62L-BV510 (BioLegend, 104441, clone MEL-14; dilution 1:200), CD44-BV605 (Biolegend, 103047, clone IM7; dilution 1:100), CD40L-PE (BD Pharmingen, 553658, clone MR1; dilution 1:100), IFNg-BV421 (BioLegend, 505830, clone XMG1.2; dilution 1:100), IL-17a-FITC (invitrogen, 11-7177-81, clone eBio17B7, dilution 1:100), FOXP3-PE (invitrogen, 12-5773-82, clone FJK-16s; dilution 1:100), and Aqua Fluorescent Reactive Dye (invitrogen, L34957 A)

### Electrocardiography

Mice were anesthetized at 2% isoflurane in accordance with IACUC protocol and one-lead electrocardiogram (lead pins inserted in right arm and left leg) was performed using AD Instruments Power Lab-C and continuous ECG tracing was recorded for 30 seconds and analyzed with LabChart Pro 8. Arrhythmias were defined as high grade atrioventricular block or ventricular arrhythmias including premature ventricular contractions and ventricular escape rhythms.

### Histology and Lesional Area analysis

Organs were dissected from mice and fixed in 10% formalin for 48h and then transferred to 70% ethanol. Samples were grossed, embedded, sectioned and H&E stained at AML Laboratories. Inflamed myocardium was quantified on H&E whole-slide images using a custom Python pipeline (**Figure S5)**. Ground-truth patches were sampled from slides of mice with confirmed myocarditis and healthy controls, manually labeled by a trained observer, and split into training and held-out test sets. Slides were color-normalized and decomposed into hematoxylin and eosin channels by stain deconvolution. From each patch, a feature vector combining stain intensity, texture (local binary patterns and gray-level co-occurrence statistics), and nuclear density was extracted. A Random Forest classifier was trained with stratified cross-validation and data augmentation, achieving ROC AUC ≥ 0.99 for inflammation detection in both cross-validation and the held-out test set. For whole-slide inference, patches were classified across the slide, the resulting probability map was upsampled and thresholded, and inflammation was reported as the percentage of tissue area above threshold. Tissue area was defined by color-space masking to exclude glass and red blood cells.

### Immunofluorescence

Harvested heart tissue was embedded in OCT, flash-frozen, and sectioned at 8 μm. Sections were fixed in either −20°C acetone (CD8/CD4/DMD), 10% NBF (cleaved-caspase-3/perforin), or 5% NBF followed by −20°C acetone (cleaved-caspase-3/CD8). For cleaved caspase-3 and perforin staining, sections were permeabilized in 0.1% NP-40 for 15 min. All sections were then blocked for 60 min in Protein Block (Abcam, ab64226). Primary antibodies ([Table S1]) were applied overnight at 4°C, followed by secondaries for 1h at room temperature, with PBS/PBS-T washes between steps. Nuclei were counterstained with DAPI and slides mounted with ProLong Gold (Life Technologies, P36930). Images were acquired on a Zeiss Axio Examiner Z1 with a 10x objective. Cleaved-caspase-3 area fraction was quantified using a custom Fiji (ImageJ) macro. Whole-tissue ROI was defined from heavily blurred (σ = 50), Otsu-thresholded DAPI images. The cleaved-caspase-3 area paired channel was background-subtracted using a rolling ball and thresholded at a fixed intensity value derived from wildtype controls to yield near-zero signal and applied identically across all samples. Cleaved-caspase-3 staining within the tissue ROI were segmented by particle analysis and its area was expressed as a fraction of total tissue area.

### Antibody Treatment

At 21 days of age, *Lag3^-/-^,Pdcd1^-/-^* mice were randomly assigned to anti-CD4 antibody (BioXCell, BE00003-1), anti-CD8 antibody (BioXCell, BE0061), anti-CD40L antibody (BioXCell, BE0017-1), anti-FasL antibody (BioXCell, BE0319), rat IgG1 isotype control (BioXCell, BE0088) or Armenian hamster IgG isotype control (BioXCell, BE0091) administered at a dose of 250ug i.p. three times weekly in 100uL. Experiments were concluded when mice reached 7 weeks of treatment. Weekly EKGs were performed until time of death or experimental endpoint. Organs were harvested for histology. To detect an anticipated mortality difference of 100% (for control) to 40% (for an intervention that reduces mortality) with an α of 0.05 and 80% power, a sample size of 8 mice per group was used.

### Statistical Analysis

Mouse survival was assessed via Kaplan-Meier Survival curves with log rank statistical analysis. Significance of quantification results was tested by Welch’s t test, Student’s t test or Mann-Whitney test using Prism 10.0 (GraphPad Software, San Diego, CA). P value <0.05 was considered as significant. Overlaid bar graphs show mean and error bars show standard error measurement.

### Study Approval

All mouse experiments were performed in accordance with an IACUC protocol (AN207188).

## Data Availability

Cardiac CD4^+^ and CD45^+^ sorted scRNA-seq data and all other data generated in this study are available on Dryad for peer review (DOI: 10.5061/dryad.7sqv9s585).

Published scRNA-seq data from patients with ICI-associated myocarditis are available in the Gene Expression Omnibus (GSE228597).

## Acknowledgements

The authors thank Sarah Elmes, Danielle Peterson, and the University of California, San Francisco (UCSF) Helen Diller Family Comprehensive Cancer Center Laboratory for Cell Analysis for use of flow cytometers and cell sorters. They thank Mylinh Bernardi and Felicia Miller of the Gladstone Genomics Core for their assistance with scRNA-seq library preparation. Sequencing was performed at the UCSF Center for Advanced Technology, supported by UCSF Program in Breakthrough Biomedical Research, Research Resource Program Institutional Matching Instrumentation Awards, and the National Institutes of Health (grant No. 1S10OD028511-01).

## Funding

American Heart Association grant 26CDA1592374 (AZM)

Sarnoff Foundation Scholar Career Development Award (AZM)

Chan Zuckerberg Biohub SF PSTP (AZM)

National Institutes of Health grant K08HL181180 (AZM)

National Institutes of Health grant R01HL15590 (JJM)

National Institutes of Health grant R01HL156021 (JJM)

National Institutes of Health grant R01HL160688 (JJM)

National Institutes of Health grant R01HL170038 (JJM)

National Institutes of Health grant P01HL141084 (JJM)

## Author contributions

Conceptualization: AZM, AG, CJK, JJM

Methodology: AZM, AG, CJK, RG, RAB

Investigation: AZM, AG, CJK, RG, RAB, ACS, ER, EO, AN

Visualization: AZM, AG, CJK, RG, RAB

Funding acquisition: JJM

Project administration: JJM

Supervision: AZM, JJM

Writing – original draft: AZM

Writing – review & editing: AG, CJK, RG, RAB, JJM

The three co-first authors contributed comparably to the study; author order was determined by the relative scope of their contributions to study conception, experimental execution, data analysis, and manuscript preparation.

## Supplementary Materials

## Supplemental Figures

**Supplemental Figure S1.**
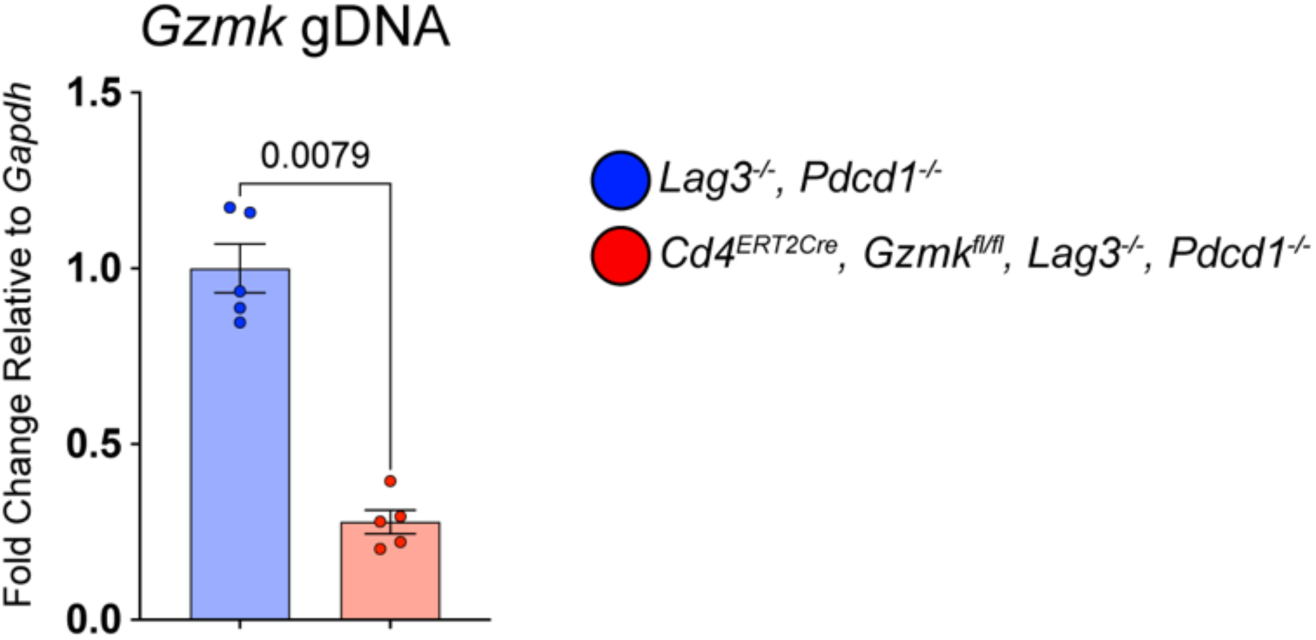
Verification of *Gzmk* DNA excision in tamoxifen-treated *Cd4-*Cre^ERT2^ *,Gzmk^fl/fl^,Lag3^-/-^,Pdcd1^-/-^* mice

**Supplemental Figure S2.**
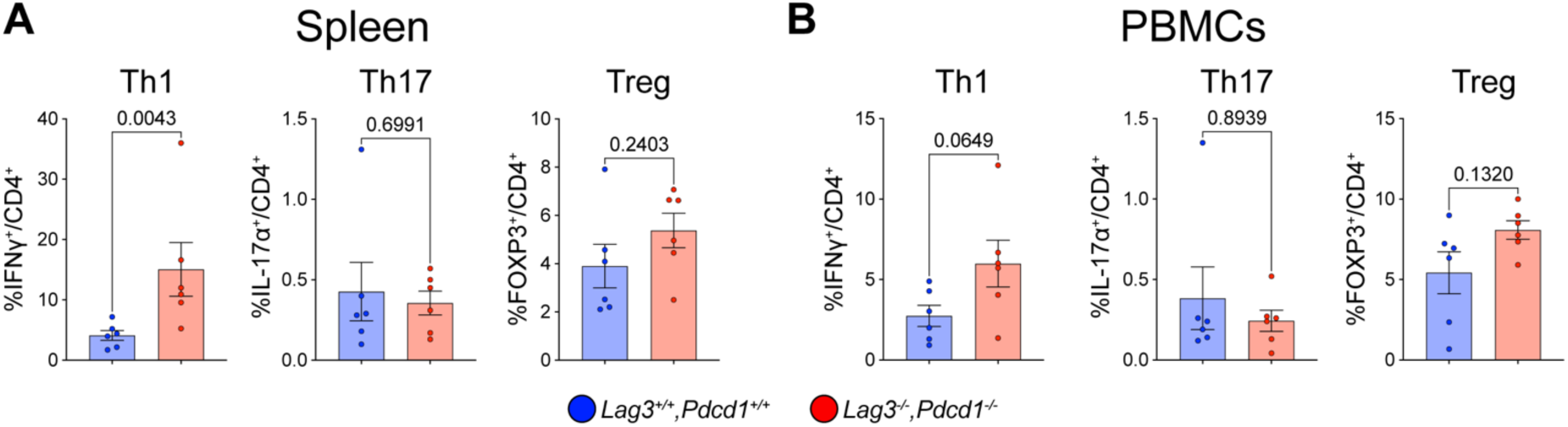
**A.** CD4^+^ helper T-cell subset flow in the spleens of *Lag3^-/-^,Pdcd1^-/-^* and *Lag3^+/+^,Pdcd1^+/+^* mice. **B.** CD4^+^ helper T-cell subset flow in the PBMCs of *Lag3^-/-^,Pdcd1^-/-^* and *Lag3^+/+^,Pdcd1^+/+^* mice.

**Supplemental Figure S3.**
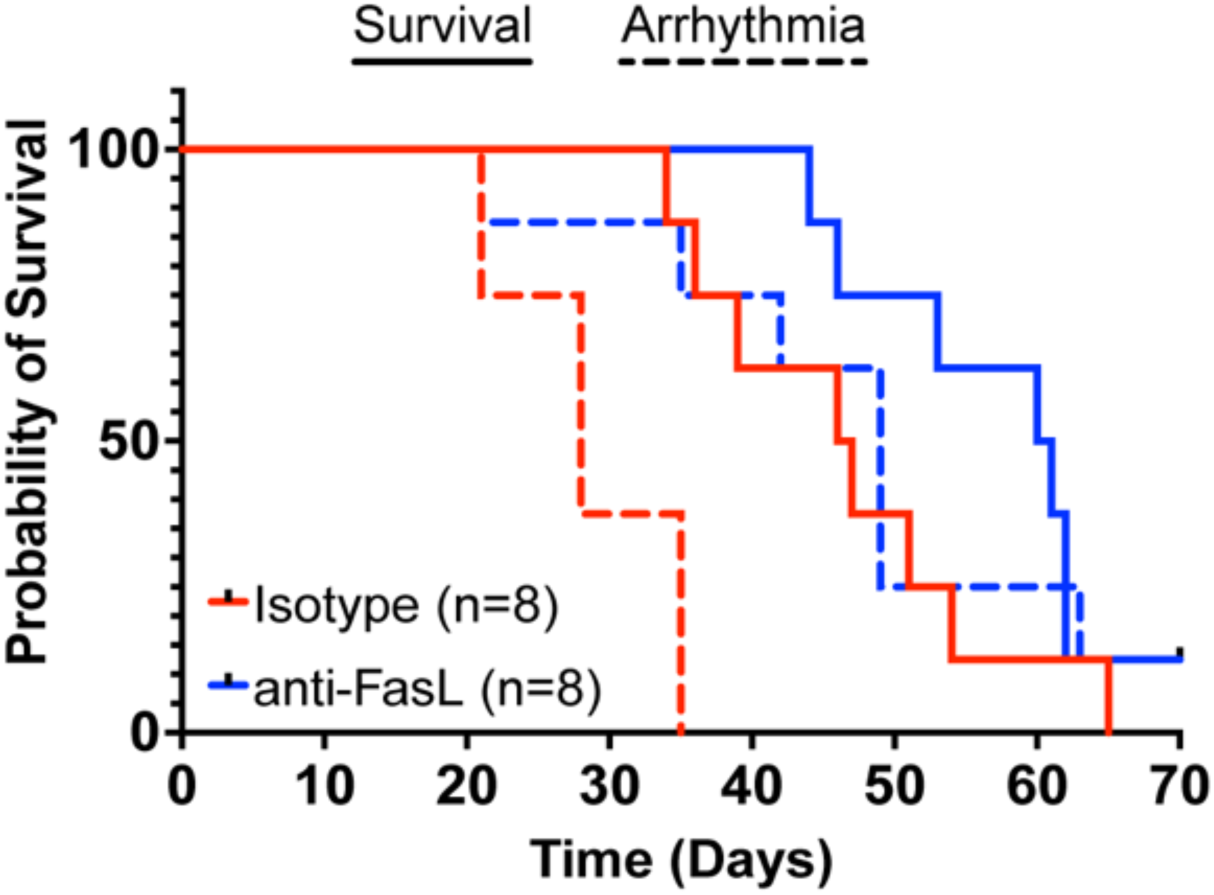
Survival curve of Isotype and anti-FASL treatment in *Lag3^-/-^,Pdcd1^-/-^* mice.

**Supplemental Figure S4.**
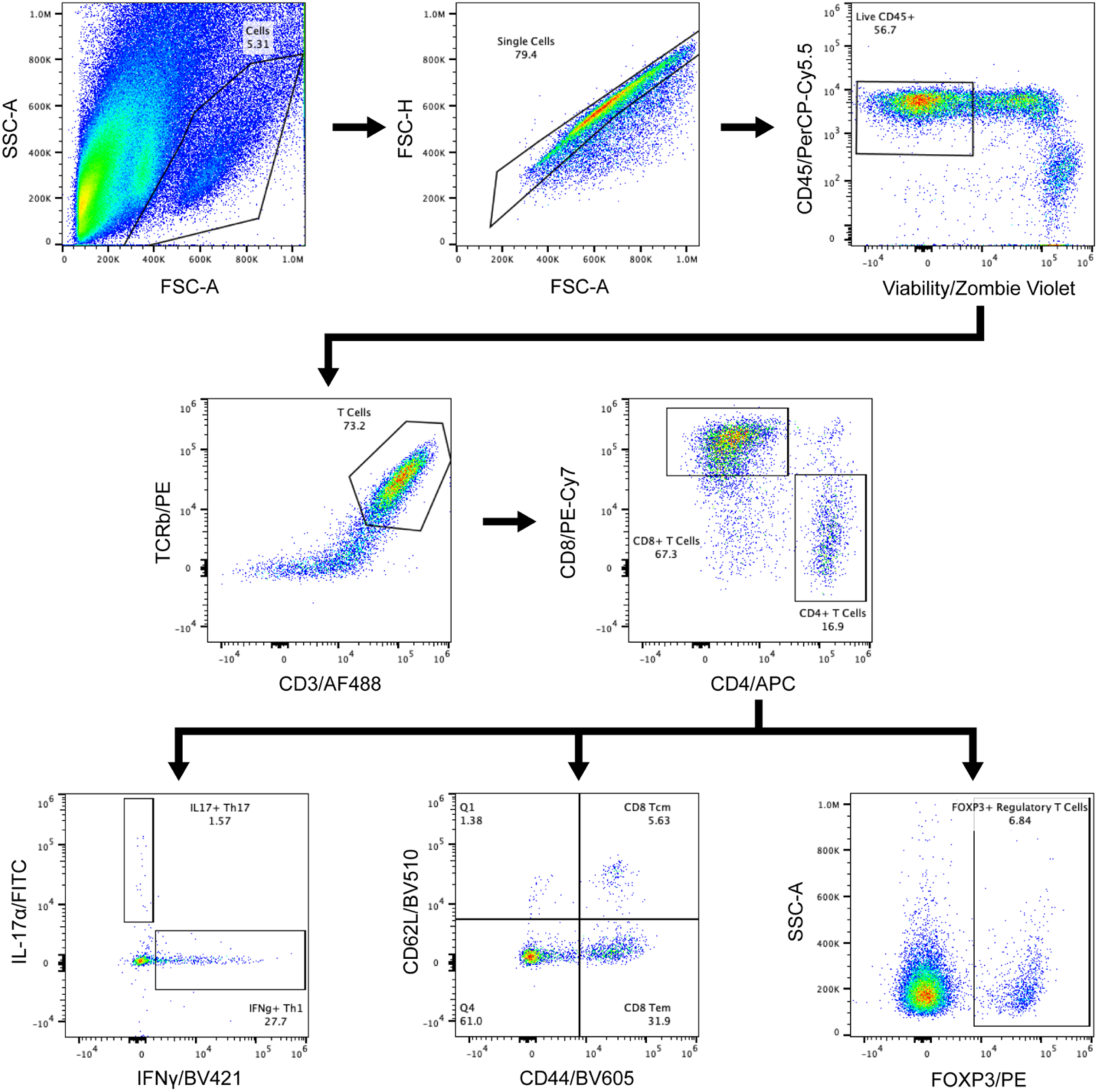
Representative gating strategy of immune populations for flow cytometry.

**Supplemental Figure S5.**
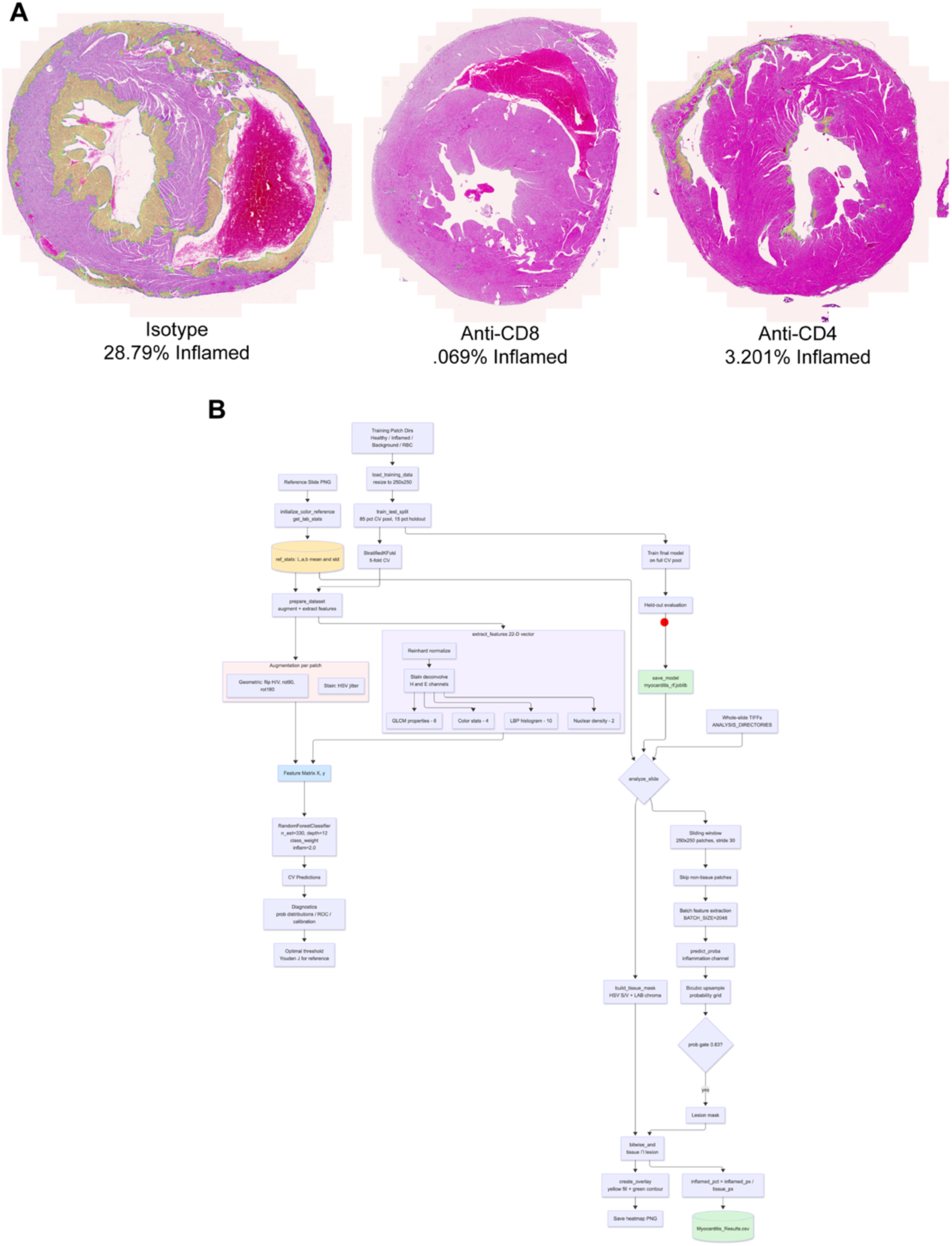
**A.** Representative H&E-stained cardiac sections from isotype control, anti-CD8, and anti-CD4-treated mice processed through the lesional area quantification pipeline. Yellow overlays demarcate regions classified as high-confidence inflammatory infiltrate. **B.** Schematic illustrating the computational workflow of the lesional area quantification pipeline.

**Supplemental Table 1.**

| Target / Reagent | Cat. No. | Manufacturer | Host | Isotype | Reactivity | Conjugate / Detection |
| --- | --- | --- | --- | --- | --- | --- |
| Primary antibodies & directly conjugated stains |  |  |  |  |  |  |
| Cleaved Caspase-3 (Asp175) | 9661T | Cell Signaling Technology | Rabbit | IgG | Human, Mouse, Rat |  |
| Dystrophin | ab15277 | Abcam | Rabbit | IgG | Human, Mouse, Rat | Unconjugated |
| CD8α | MA1-10301 | Invitrogen | Rat | IgG2a, κ | Mouse | Unconjugated |
| CD4 | 14-0041-82 | Invitrogen (eBioscience) | Rat | IgG2b, κ | Mouse | Unconjugated |
| DAPI (nuclear stain) | 62248 | Invitrogen | N/A | N/A | Universal | DAPI |
| WGA (wheat germ agglutinin) | W11261 | Invitrogen (Molecular Probes) | N/A | N/A | Universal | Alexa Fluor 488 |
| Secondary antibodies |  |  |  |  |  |  |
| Goat anti-Rat IgG (H+L) | A21434 | Invitrogen | Goat | IgG | Rat IgG (H+L) | Alexa Fluor 555 |
| Donkey anti-Rabbit IgG (H+L) † | A31570 | Invitrogen | Donkey | IgG | Rabbit IgG (H+L) | Alexa Fluor 647 |

